# SlSAURs, targeted by SlSTOP1, inhibit SlPP2C.Ds to modulate H^+^-ATPase and tomato root elongation under aluminum stress

**DOI:** 10.1101/2024.05.10.593632

**Authors:** Danhui Dong, Qilin Deng, Jialong Zhang, Congyang Jia, Lei Zhang, Hongxin Li, Na Zhang, Yang-Dong Guo

## Abstract

Aluminum (Al) stress, a prevalent constraint in acid soils, is known to inhibit plant growth by inhibiting root elongation through restricted cell expansion. The molecular mechanisms of Al-induced root inhibition, however, are not fully understood. This study aimed to elucidate the role of SlSAUR (Small auxin up-regulated RNA) proteins, which were downstream of the key Al stress-responsive transcription factor SlSTOP1 and its enhancer SlSZP1, in modulating root elongation under Al stress in tomato (*Solanum lycopersicum*). Our findings demonstrated that tomato lines with *SlSAURs* knockout exhibited shorter root lengths when subjected to Al stress. Further investigation into the underlying mechanisms revealed that SlSAURs interact with D-clade type 2C protein phosphatases, specifically SlPP2C.Ds. This interaction was pivotal as it suppresses the phosphatase activity, leading to the derepression of SlPP2C.D’s inhibitory effect on H^+^-ATPase. Consequently, this promoted cell expansion and root elongation under Al stress conditions. Our research significantly contributes to the current understanding of the molecular mechanisms by which Al ions modulate root elongation. The discovery of the SlSAUR-SlPP2C.D interaction and its impact on H^+^-ATPase activity provides a novel perspective on the adaptive strategies employed by plants to cope with Al toxicity. This knowledge may pave the way for the development of tomato cultivars with enhanced Al stress tolerance, thereby improving crop productivity in acid soils.

## Introduction

Aluminum is the third most abundant element in the crust, accounting for approximately 8% by weight and ranks first among all metallic elements. However, the abundant aluminum can cause toxicity to plants. When the soil pH falls below 5.5, the aluminum, which originally existed in a non-toxic form, such as aluminum oxides and aluminum silicates, will be dissolved into free trivalent aluminum ions (Al^3+^). These ions are highly deleterious to plant growth and development, with the most notable impact inhibiting the growth of primary root elongation (Ownby and Popham, 1990). Even brief exposure to Al-containing solutions can inhibit root growth (Llugany et al., 2008).

STOP1 (Sensitive to Proton Rhizotoxicity 1), a C2H2 zinc-finger transcription factor, is a central regulator of stress regulation (Sadhukhan et al., 2021), especially in adapting to acid soil (Iuchi et al., 2007). The *stop1* demonstrated heightened sensitivity to protons and an even greater sensitivity to aluminum, resulting in markedly inhibited primary root elongation under conditions of aluminum stress (Iuchi et al., 2007). In *stop1*, key genes associated with aluminum tolerance, such as *ALMT1* and *ALS3*, were significantly downregulated (Sawaki et al., 2009). Additionally, the *POLYGALACTURONASE-INHIBITING PROTEIN 1* (*PGIP1*), which stabilized pectin in the cell wall under acid soil, was regulated by STOP1 (Agrahari et al., 2021). Furthermore, the *GLUTAMATE-DEHYDROGENASE 1* and *2* (*GDH1* and *GDH2*), identified as novel Al resistance genes, were also regulated by STOP1 and involved in Al tolerance (Tokizawa et al., 2021).

Current research has primarily focused on the downstream target genes of STOP1 and their roles in mitigating Al toxicity by mechanisms such as organic acid exudation, aluminum ion transport, and regulation of cellular pH homeostasis and metabolic processes. However, the direct regulation of root elongation by these proteins remains unclear. The fundamental mechanisms underlying Al-induced inhibition of root elongation and the identity of functional proteins that influence this process are areas that warrant further investigation.

The acid growth hypothesis, which proposes that auxin stimulates cell expansion by acidifying the cell wall, provides a framework for understanding how *small auxin up-regulated RNAs* (*SAURs*) might be involved in this process. *SAUR* was first identified in soybean as rapidly responding to auxin (Mcclure and Guilfoyle, 1987). Overexpression of certain *SAUR* genes, such as *SAUR63*, has been linked to increased cell elongation, as observed in hypocotyls, petals, and stamen filaments (Chae et al., 2012). The SAUR19 subfamily, including SAUR19-24, was associated with larger leaf size and increased hypocotyl elongation (Spartz et al., 2012).

Phosphorylation at Thr947 was significant for activation of plasma membrane (PM) H^+^-ATPases, which is integral to the acid growth hypothesis (Takahashi et al., 2012). The overexpression of *SAUR19* resulted in an increased PM H^+^-ATPase activity and reduced apoplastic pH (Spartz et al., 2014; Fendrych et al., 2016). This overexpression also resulted in an increase in the phosphorylation level of Thr947 on AHA2, which consistent with the elevated AHA Thr947 phosphorylation when indole-3-acetic acid (IAA) or fungal toxin fusicoccin (FC) were applied (Spartz et al., 2014). SAUR19 and other SAURs have been demonstrated to interact with a subset of type 2C protein phosphatases known as D-clade PP2Cs (PP2C.D), which can dephosphorylate Thr947 on AHA2 (Spartz et al., 2014; Ren et al., 2018; Wong et al., 2019). This interaction at the plasma membrane is crucial as it inhibits the activity of PP2C.D phosphatases, thereby potentiating the activity of PM H^+^-ATPases (Spartz et al., 2014; Wang et al., 2020; Cui et al., 2023), and facilitating cell expansion. Collectively, these findings consolidate the genetic evidence supporting the acid growth theory and refine the mechanistic understanding of how SAURs, in conjunction with PP2C.D and PM H^+^-ATPases, orchestrate cellular expansion in response to auxin signaling.

Despite the growing evidence linking SAURs to cell expansion, research on their broader role in plant development and stress responses remains limited. The regulatory elements acting upstream of SAURs are also not well characterized. It was discovered that the ARF-BZR-PIF complex co-regulated the expression of *SAUR* to promote hypocotyl elongation (Oh et al., 2014), and individual TF (transcription factor) of the complex could regulate *SAUR* expression in plant growth and development. For instance, ARF7/19, binding to the promoter of *SAUR19*, promoted the gravitropism and phototropism of plant hypocotyls (Wang et al., 2020). TCP4 and PIF3 antagonistically regulated *SAURs*, playing a role in cotyledon opening and closing (Dong et al., 2019). The zinc-finger proteins AZF1 and AZF2 have been reported to suppress *SAUR* genes, negatively impacting plant growth under abiotic stress conditions (Kodaira et al., 2011). Despite the recognized importance of SAURs in plant development and stress responses, including their induction by abscisic acid (ABA) and drought treatment (He et al., 2021), the specific TFs within regulatory complexes that act upstream of SAURs require further investigation. This is particularly relevant given the diverse membership of SAURs across different species and their induction under various stress conditions, such as the saline-alkali stress-induced TaSAUR215 in wheat, which interacted with TaCCD1 to enhance stress tolerance (Cui et al., 2023), and the *MdSAUR-like* may have connection to fruit ripening in apple (Wang et al., 2024).

Tomato, as one of the most produced vegetables in the world, are increasingly affected by soil acidification, a condition exacerbated by excessive fertilizer application in facility cultivation (Guo et al., 2010). The mobility of aluminum ions in soil caused by soil acidification poses a serious threat to plant growth. Our previous study has revealed that SlSTOP1, enhanced by SlSZP1, plays a crucial role in tomato Al tolerance, exhibits inhibited root elongation under Al stress when loss of function mutated (Zhang et al., 2022). While the underlying cause of root growth inhibition in *Slstop1* mutants under Al stress remains elusive. Given that aluminum mainly reduced cell expansion of root (Doncheva et al., 2005), identifying the fundamental factors that regulate cell expansion under Al stress is a critical research question.

In this study, we have identified two genes, *SlSAUR36* and *SlSAUR38*, which are downstream of SlSTOP1 and its enhancer SlSZP1.These proteins have been shown to interact with the protein phosphatase SlPP2C.D. This interaction reduces the activity of phosphatase to release the inhibition of SlPP2C.D to PM H^+^-ATPase SlLHA4. This modulation activates the H^+^-ATPase, thereby promoting the cell expansion. Our findings provide novel insights into the regulatory mechanism governing primary root length under Al stress mediated by SlSTOP1. This study aims to elucidate the molecular dialogue between Al toxicity, root elongation, and the regulatory network involving SlSAURs, offering a comprehensive perspective on how plants adapt to and mitigate the effects of Al stress.

## Results

### SlSTOP1 positively regulates *SlSAUR36* and *SlSAUR38* expression

In previous studies, we observed a significant inhibition in root elongation of *Slstop1* under aluminum stress (Zhang et al., 2022). Through an analysis of RNA-seq data from *Slstop1* under aluminum stress, we identified two *SlSAURs* responding to aluminum ions and were downregulated by *Slstop1* knockout (Figure S1A). *SlSAUR36 and SlSAUR38* showed a significant increase in mRNA levels-approximately 50-fold for *SlSAUR36* and 40-fold for *SlSAUR38* under aluminum stress, indicating a dependency on SlSTOP1 for their expression. Therefore, we speculate that they may function downstream of SlSTOP1 in regulating root elongation under aluminum stress.

To explore this further, we tested the expression dynamics of *SlSAUR36* and *SlSAUR38* in Ailsa Craig tomato under 60 μM aluminum ions at various time points. In wild type plant, after 9-hour aluminum treatment, the mRNA levels of *SlSAUR36* and *SlSAUR38* were increased by 15 and 20 times (Figure S1B and S1C), indicating that *SlSAUR36/38* may respond to aluminum stress subsequent to the accumulation of SlSTOP1. Subsequent RT-qPCR positively validated the transcriptome data, showing that under aluminum stress, *SlSAUR36* and *SlSAUR38* were significantly induced, with an upregulation of about 20-fold in wild type. Notably, their expressions were significantly inhibited in the *Slstop1* mutant under aluminum ions containing conditions, with downregulation multiples of nearly 5 for *SlSAUR36* and 20 times for *SlSAUR38*, respectively (Figure 1A and 1B).

**Figure 1.**
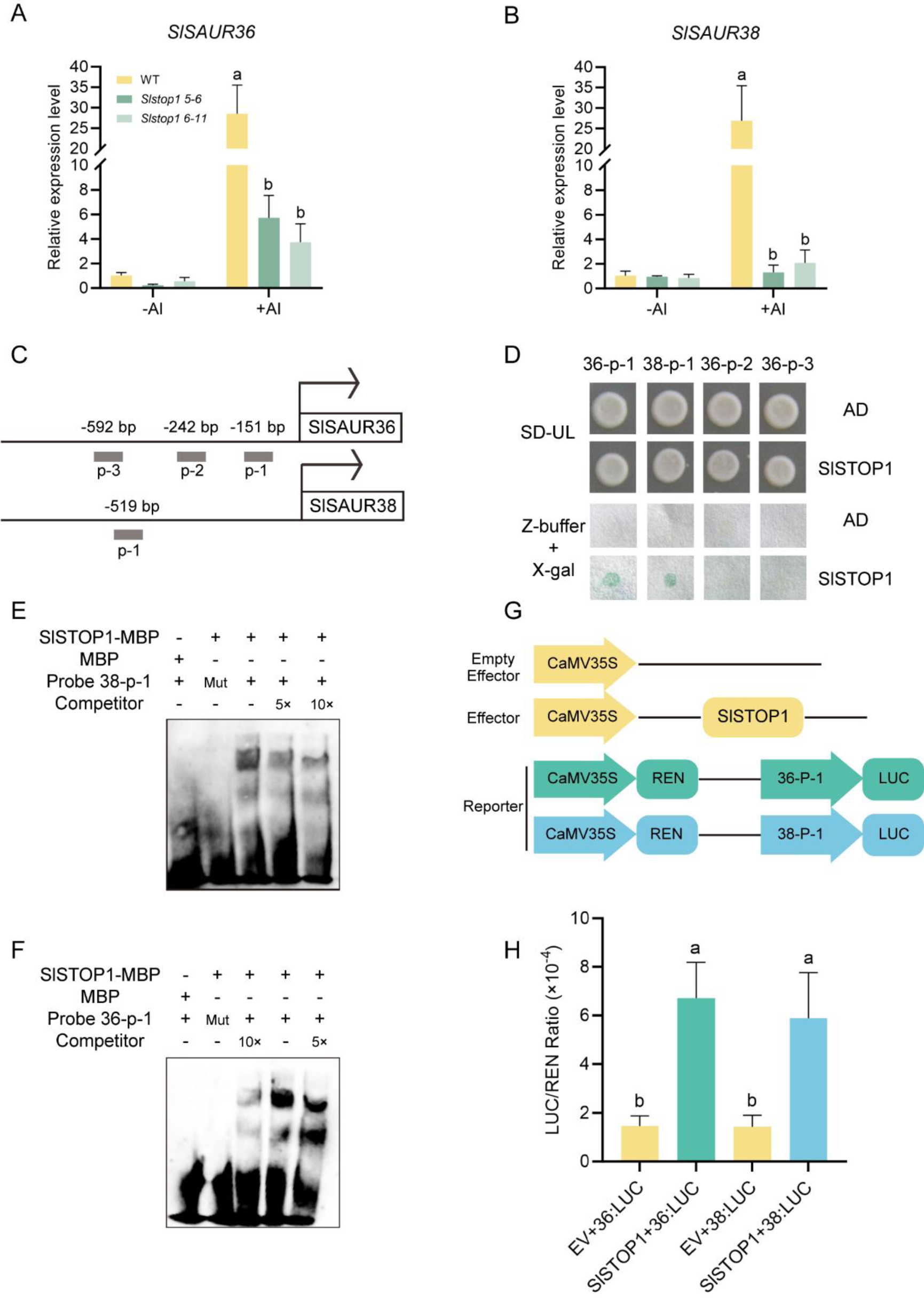
SlSTOP1 targets *SlSAUR36/38* to positively regulates their expression. **A and B** The relative expression level of *SlSAUR36* and *SlSAUR38* in wild type and *Slstop1* with or without treated by 60 μM AlCl_3_ (pH 4.7). Data were presented as means ± SD (n=3). Different lowercase letters indicate significantly different means (P < 0.05, ANOVA test followed by Tukey test). **C and D** Y1H assay. P-s indicated the fragment containing GGNVS or cis-D of *SlSAUR36* and *SlSAUR38*. The fragment was cloned into pLacZi vector, SlSTOP1 was cloned into pGAD424 vector. Transformants with paired constructs were grown on SD-UL medium and then used for binding assay in Z-buffer with X-gal. **E and F** EMSA assay. Two p-1s were labelled with or without biotin, and corresponding mutant probe was also labelled with biotin. The probe and SlSTOP1-GST were co-incubated for binding assay. **G** Schematic diagrams of effector and reporter constructs used for LUC/REN assay. **H** LUC/REN assay. SlSTOP1 was cloned into pGreen II 62-SK as the effector. The p-1 of *SlSAUR36* and *SlSAUR38* were fused with the LUC as reporters, respectively. Empty vector (pGreen II 62-SK) co-expressing with reporter was set as the control. Data were presented as means ± SD (n=10). Different lowercase letters indicated significantly different means (P < 0.05, ANOVA test followed by Tukey test).

STOP1 and its homolog OsART1 have been shown to regulate the expression of downstream target genes by binding to specific cis-elements, such as D and GGNVS motifs (Tsutsui et al., 2011; Tokizawa et al., 2015). By analyzing the 1500 bp upstream region of the promoter for *SlSAUR36* and *SlSAUR38*, we found some potential binding sites for these elements, which were designated as p-1, p-2, and p-3 for further analysis (Figure 1C). We then validated the in vitro binding of SlSTOP1 to these segments using yeast one hybridization. The results showed that SlSTOP1 could bind to the p-1 segment of both *SlSAUR36* and *SlSAUR38* (Figure 1D). The binding was further validated by electrophoretic mobility shift assays (EMSA), which demonstrated that a GST-tagged SlSTOP1 protein, purified from *E. coli*, could bind to biotin-labeled probes (Figure 1E and 1F). Next, we used tobacco for LUC/REN assay to validate transcriptional activation of SlSTOP1 to *SlSAUR36* and *SlSAUR38* in vivo. The addition of SlSTOP1 to the system resulted in a significant increase in the LUC/REN ratio, indicative of transcriptional activation (Figure 1G and 1H). These results demonstrate that SlSTOP1 can bind to promoters of *SlSAUR36* and *SlSAUR38*, thereby positively regulating their expression.

### SlSTOP1 and SlSZP1 synergize to regulate *SlSAUR36/38* expression under aluminum stress

Our previous findings indicated that SlSZP1 interact with SlSTOP1, thereby improving the stability of SlSTOP1 protein (Zhang et al., 2022). Under aluminum stress, *Slszp1* mutant, similar to the *Slstop1* mutant, showed shorter root length compared to WT (Zhang et al., 2022). RNA-seq of *Slszp1* also suggested that SlSTOP1 and SlSZP1 may operate within the same pathway to confer aluminum resistance. This led us to hypothesize that SlSZP1 might also regulate the expression of *SlSAUR36* and *SlSAUR38*, contributing to the plant’s response to aluminum stress. To test this hypothesis, we screened members of the *SlSAUR* family from the *Slszp1* RNA-seq and analyzed their expression patterns. Notably, we found the expression of *SlSAUR36* and *SlSAUR38* was significantly repressed, approximately 20-fold and 80-fold, respectively, in the absence of SlSZP1 (Figure 2A). This promoted us to investigate whether SlSZP1 could bind to the promoters of *SlSAUR36* and *SlSAUR38* to modulate their expression. Given that SlSZP1 is also a C2H2 type TF, we still selected the p-1, p-2 and p-3 sites as the potential binding region. Y1H assay preliminarily validated these binding (Figure 2B and 2C), confirming that SlSZP1 could bind to p-1 of both *SlSAUR36* and *SlSAUR38*. We further used LUC/REN to assess the transcriptional activation of SlSZP1 to *SlSAUR36* and *SlSAUR38* in vivo (Figure 2D and 2E). The addition of SlSZP1 led to a significant increase in the LUC/REN ratio, confirming that SlSZP1 can activate the expression of *SlSAUR36* and *SlSAUR38*. These results indicate SlSZP1 not only promotes the accumulation of SlSTOP1 at the protein level (Zhang et al., 2022), thereby increasing the expression of downstream genes, but also directly binds to the same site of *SlSAURs* to promote the expression, albeit with a potential temporal delay compared to SlSTOP1-mediated activation. Moreover, when we co-expressed SlSTOP1 and SlSZP1, we observed a higher increase in the LUC/REN ratio than with SlSZP1 alone (Figure 2D and 2E). This indicates that in tomato plants, SlSTOP1 and SlSZP1 may function synergistically to regulate the expression of *SlSAUR36* and *SlSAUR38* during aluminum toxicity. Additionally, each factor can operate independently, exhibiting transcriptional activity at different time points throughout the aluminum stress response.

**Figure 2.**
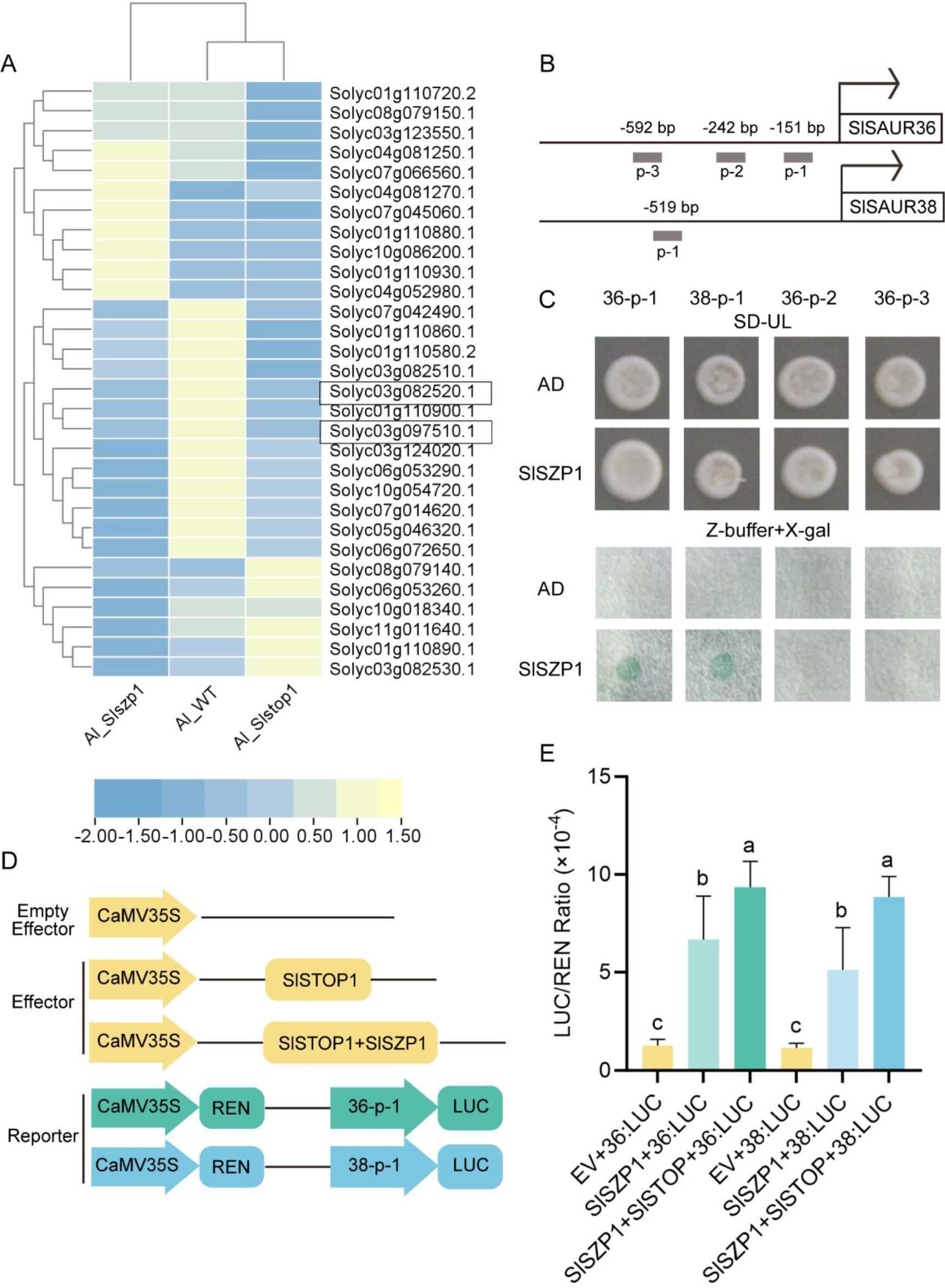
SlSTOP1 and SlSZP1 synergistically enhance the expression of *SlSAUR36/38*. **A** Heatmap representation of the expression analysis for *SlSAUR* genes in *Slstop1*, *Slszp1* and WT treated with 60 μM AlCl_3_ for 9 hours (pH 4.7). The RNA-sequencing data have been deposited in the Gene Expression Omnibus (GEO) database under the accession number GSE168433 and GSE201111. The heatmap illustrated the mean expression levels ± SD across three biological replicates (n=3). **B and C** Y1H assay. P-s indicated the fragment containing GGNVS or cis-D of *SlSAUR36* and *SlSAUR38*. The fragment was cloned into pLacZi vector, SlSZP1 was cloned into pGAD424 vector. Transformants with paired constructs were grown on SD-UL medium and then used for binding assay in Z-buffer with X-gal. **D** Schematic diagrams of effector and reporter constructs used for LUC/REN assay. **E** SlSTOP1 and SlSZP1 were cloned into pGreen II 62-SK respectively as the effector. Empty vector (pGreen II 62-SK) co-expressing with reporter was set as the control. Data were presented as means ± SD (n=10). Different lowercase letters indicated significantly different means (P < 0.05, ANOVA test followed by Tukey test).

### *SlSAUR36* and *SlSAUR38* function redundantly in root elongation under aluminum stress

SlSAUR36 and SlSAUR38 belong to the auxin inducible superfamily (Figure S2A). Tissue expression analysis revealed that both *SlSAURs* have expression at root, especially *SlSAUR38*, which is specifically expressed in root (Figure S2B). Multiple sequence alignment analysis suggests significant variation at the N terminal and C terminal, but the SAUR family domain remains conserved (Figure S3A). Studies indicated that SAURs localized to distinct cellular compartments to perform various functions. Predictively, the molecular weight of SlSAUR36 and SlSAUR38 were determined to be 14 and 18 kDa, respectively. It is well established that proteins smaller than 40 kDa can diffuse in and out of the nucleus (Marathe et al., 2024). To accurately determine the subcellular locations of these proteins, we fused two GFP tags with SlSAURs to enlarge the proteins and prevent diffusion into the nucleus. The result showed that both SlSAUR36 and SlSAUR38 were located in nuclear, cytoplasm and plasma membrane (Figure S3B). Given that MeSAUR1 functioned as a transcription factor in *cassava* (Ma et al., 2017), we tested whether SlSAUR36/38 could also function as transcription factors refer to their nucleus localization. When the full lengths of SlSAUR36 and SlSAUR38 were fused with GAL4-BD, no transcription activation signal was observed (Figure S3C). The LUC/REN ratio of SlSAUR36-GAL4-BD and SlSAUR38-GAL4-BD in protoplasts showed no significant difference compared to GAL4-BD (Figure S3D), indicating that neither SlSAUR36 nor SlSAUR38 functions as transcription regulators.

To assess the function of SlSAUR36 and SlSAUR38, we used CRISPR/Cas9 system to generate single *Slsaur36*, *Slsaur38*, and double *Slsaur36/38* knock out lines for subsequent experiments (Figure 3A and 3B). Under normal conditions, there were no difference in root length among all of the transgenic lines (Figure 3C and 3D). However, when growth medium was supplemented with 30 μM Al^3+^ for 14 days, all knock lines exhibited shorter root length, with the double *Slsaur36/38* knock out lines *Slsaur-168* and *Slsaur-11* displaying even shorter root length compared to the single knock lines (Figure 3C and 3D), indicating functional redundancy between the two genes. The overexpression lines of *SlSAUR36* and *SlSAUR38*, driven by their native promoters, showed longer root length than wild type (Figure 3C and 3D). To further validate the function of *SlSAUR36* and *SlSAUR38* under aluminum stress, we performed separate and combined complementation of *SlSAUR36* and *SlSAUR38* in *Slstop1*. The inhibition of *Slstop1* root growth was alleviated by the complementation with *SlSARU36* and *SlSAUR38* (Figure 3E and 3F). The complementation of *SlSAURs* in the corresponding mutant further confirmed the phenotype was due to the specific *SlSAURs* mutations. When complementing *SlSAUR36*, *SlSAUR38*, and *SlSAUR36 and SlSAUR38*, the phenotypes of root length in complement lines was recovered (Figure S4B), effectively ruling out the possibility of off-target effects.

**Figure 3.**
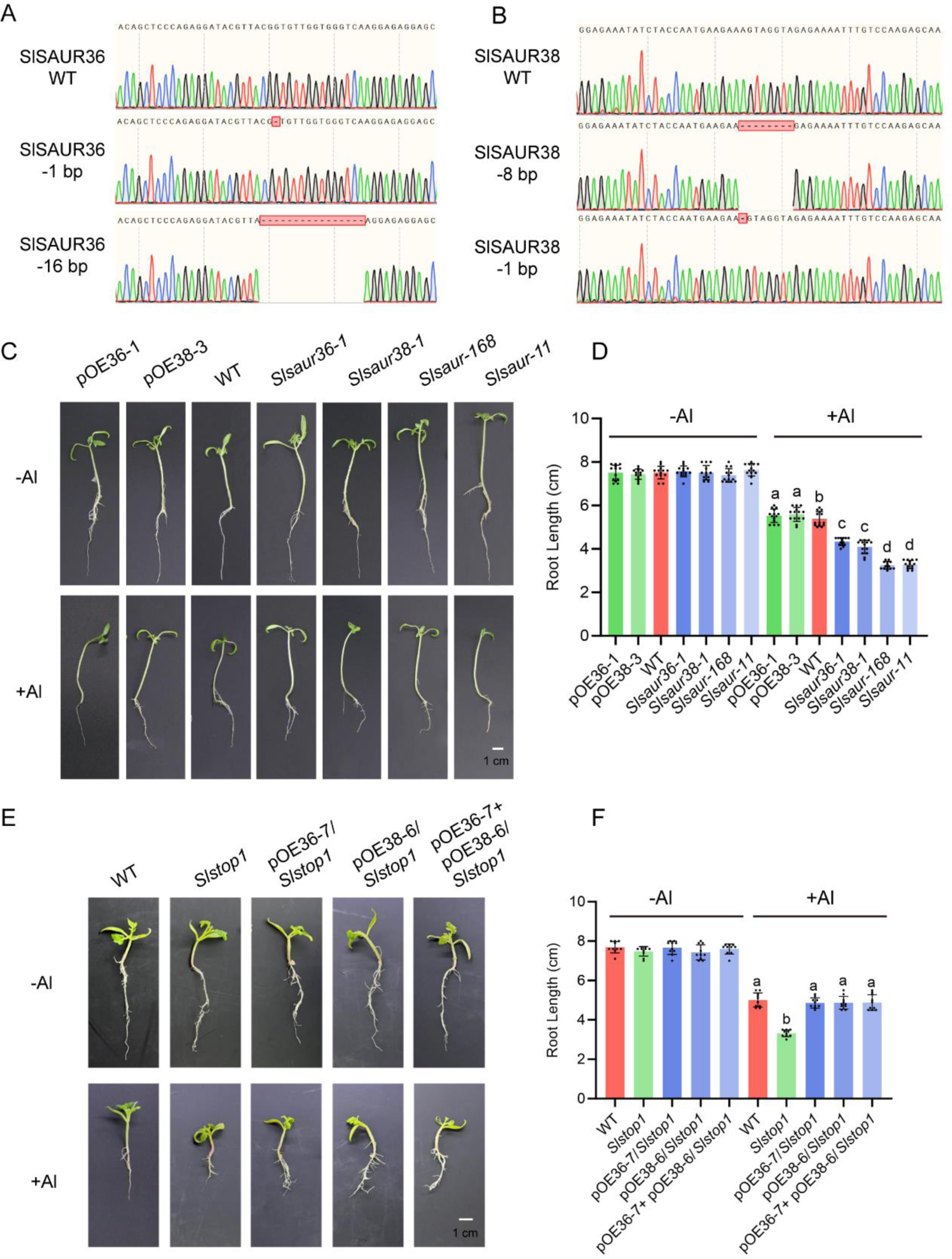
*Slsaur36* and *Slsaur38* are sensitive to aluminum stress. **A and B** Gene editing results of *SlSAUR36* and *SlSAUR38*. **C and E** Phenotype image. *Slsaur36-1* and *Slsaur38-1* were single knockout lines of *SlSAUR*, respectively. *Slsaur-168* and *Slsaur-11* were double knockout lines of *SlSAURs*. Two-week old seedlings were grown in Hoagland solution supplemented with 0 or 30 μM AlCl_3_ (pH 4.5) for 10 days. Scale bar =1 cm. **D and F** Quantitative data of C and E. Data were presented as means ± SD (n=10). Different lowercase letters indicated significantly different means (P < 0.05, ANOVA test followed by Tukey test).

### SlSAURs interact with SlPP2C.Ds to inhibit the activity of phosphatase

SAUR proteins are known to interact with D clade of PP2Cs to modulate their phosphatase activity, thereby influencing various aspects of plant biology (Wang et al., 2020; Yin et al., 2020; Cui et al., 2023). To elucidate the molecular interactions between SAUR proteins and PP2C phosphatases in the context of plant growth and development, we conducted a series of experiments. We employed a yeast two-hybrid (Y2H) system to explore the possible combinatorial interactions between SlSAUR36/38 and SlPP2C.D isoforms. Specifically, we constructed SlPP2C.Ds in a GAL4-based Y2H assay, where the SlPP2C.D was fused to the GAL4 activation domain (GAL4-AD), and SlSAUR36/38 were fused to the GAL4 DNA-binding domain (GAL4-BD). Our screening efforts identified interactions between SlSAUR36 and SlPP2C D4/D5, as well as between SlSAUR38 and SlPP2C.D2/D3 (Figure S5A). Subsequent luciferase complementation imaging (LCI) assays provided positive evidence for the interactions identified in the yeast two-hybrid system (Figure S5B). To validate and further characterize these interactions, we utilized a co-immunoprecipitation (Co-IP) approach in a tobacco system, where SlSAUR36 and SlSAUR38, each fused with a GFP tag, were shown to interact with SlPP2C.D5 and D2, respectively (Figure 4A). We also used bimolecular fluorescence complementation (BiFC) to ascertain the subcellular localization of these interactions. Our findings revealed that the interactions occur in the nucleus, cytoplasm, and plasma membrane (Figure 4B). Notably, the interactions involving SlSAUR36 with SlPP2C.D4 and SlSAUR38 with SlPP2C.D2 was not found in nucleus. To exclude false positive possibilities, we further detect the localization of the SlPP2C.D isoforms, confirming that SlPP2C.D2/3/5 were localized in nuclear, cytoplasm, and plasma membrane, while SlPP2C.D4 was only found in plasma membrane (Figure S6). Collectively, these results validate the interactions between SlSAUR proteins and SlPP2C.D phosphatases. To gain insights into the molecular basis of these interactions, we performed in silico docking analyses, which predicted specific docking sites between SlPP2C.Ds and SlSAURs (Figure 4C and S7). Furthermore, we expressed and purified SlPP2C.D as GST fusion protein and SlSAUR36/38 as MBP fusion proteins from *E. coli*. Using p-Nitrophenyl Phosphate (pNPP) as substrate, we assessed the in vitro phosphatase activity of SlPP2C.Ds. Our results demonstrated that SlPP2C.Ds exhibited measurable phosphatase activity, which was significantly reduced upon the addition of SlSAUR proteins to the reaction mixture (Figure 4D and E). In conclusion, our experimental findings establish that SlSAUR proteins can interact with SlPP2C.D phosphatases and inhibit their phosphatase activity.

**Figure 4.**
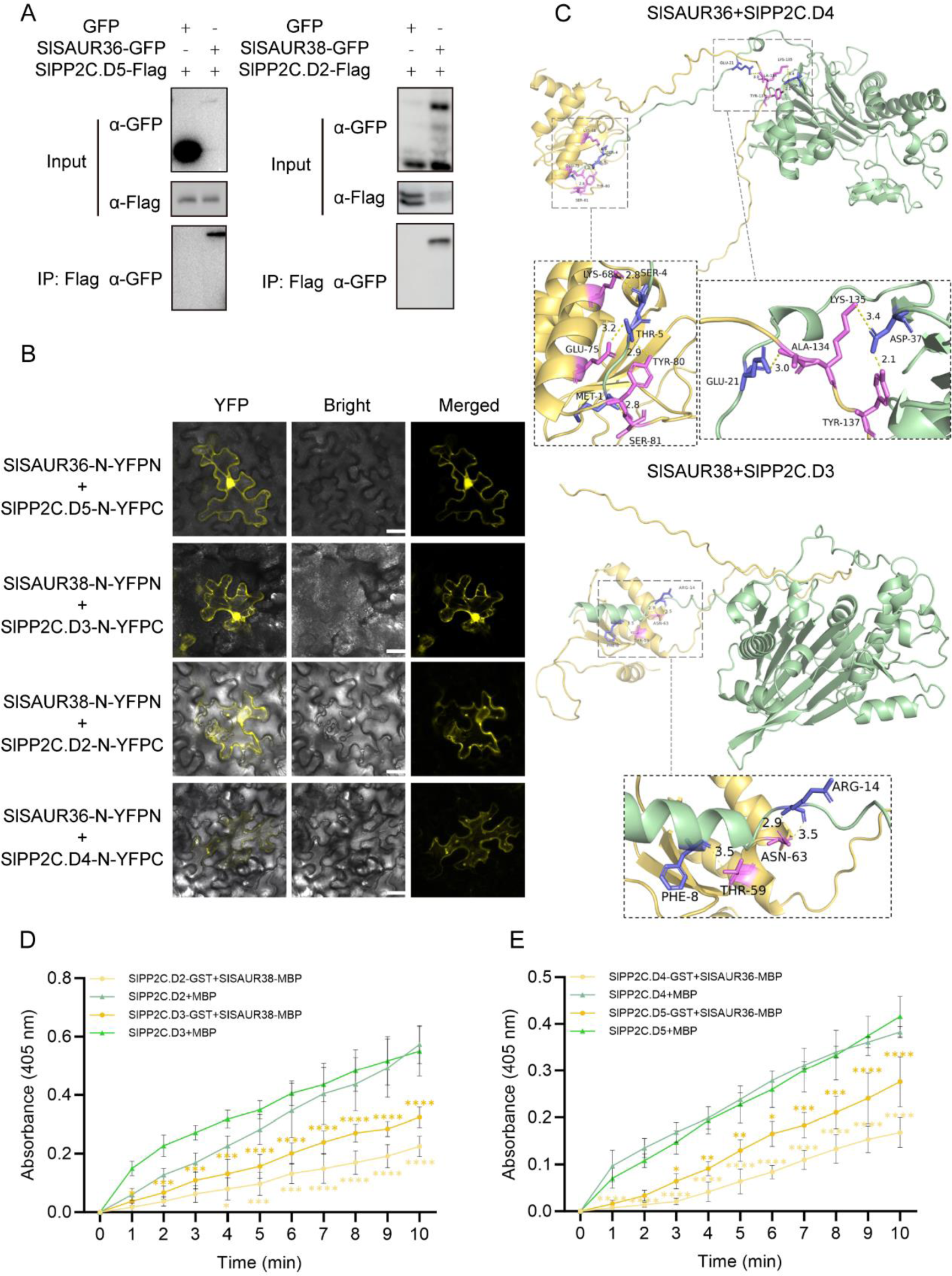
SlSAURs interact with SlPP2C.Ds to inhibit the activity of phosphatase. **A** Co-IP assay. SlSAURs-GFP was co-infiltrated with SlPP2C.Ds-Flag in N. *benthamiana* leaves. After 4 days, total protein extracts were immunoprecipitated with anti-Flag agarose beads and then detected with anti-GFP antibody. **B** BiFC assay. SlSAURs-N-YFPN and SlPP2C.Ds-N-YFPC were transiently expressed in tobacco leaves. Scale bar =50 μm. **C** Molecular docking of SlSAUR36 and SlPP2C.D4 (above), SlSAUR38 and SlPP2C.D3 (below). **D and E** SlSAURs inhibit the phosphatase activity of SlPP2C.Ds. Purified SlSAURs-MBP and SlPP2C.Ds were co-incubated in reaction buffer. SlPP2C.Ds and MBP were set as negative control. The pNPP was the substrate. Absorbance at 405 nm was recorded every minute for a total duration of 10 minutes. Data were presented as means ± SD (n=3). Asterisks indicated significant differences. (t-test, *P < 0.05, **P < 0.01, ***P < 0.001, ****, P < 0.0001).

### SlSAUR36/38 promote PM H^+^-ATPase activity by inhibiting SlPP2C.Ds phosphatase activity

PP2C.Ds are known to control cell expansion by dephosphorylating the penultimate threonine of PM H^+^-ATPases to modulate their activities (Ren et al. 2018). SAUR proteins have been shown to promote PM H^+^-ATPase activity by inhibiting PP2C.D phosphatase activity (Spartz et al., 2014). Given this context, we hypothesized that SlSAUR36/38 may regulate cell expansion in tomato through a similar mechanism involving PP2C.D and PM H^+^-ATPases. To investigate the potential regulatory role of SlSAUR36/38 in cell expansion via the PP2C.D-PM H^+^-ATPases pathway in tomato, we conducted a series of molecular and cellular biology experiments. To explore this possibility, we first constructed amino acid evolutionary tree analysis comparing plasma membrane proton pumps in *Arabidopsis* and tomato (Figure S8). In *Arabidopsis*, aluminum stress induced the expression of *AHA1/2/7* (Zhang et al., 2019), with AHA2 being the major H^+^-ATPase isoform in roots (Młodzińska et al., 2015), contributed to the root response to Al stress (Zhang et al., 2019). Through sequence alignment, we identified SlLHA4 as the closest homolog to AHA2 in tomato and selected SlLHA4 for further study. We employed BiFC, LCI, and Co-IP assays to verify the interaction on plasma membrane between SlPP2C.Ds and SlLHA4 (Figure 5A, B and C). Molecular docking analysis also supported that the physical interaction between SlLHA4 and SlPP2C.Ds (Figure S9). To assess the functional effect of the SlPP2C.Ds-SlLHA4 interaction on proton pump activity, we expressed SlLHA4 in *RS72* yeast strains, which lack endogenous H^+^-ATPase activity. We observed that yeast strains expressing *SlLHA4* were capable of growing on glucose-containing media, indicating that SlLHA4 could complement the yeast proton pump deficiency (Figure 5D). However, co-expression of SlPP2C.Ds and SlLHA4 resulted in impaired growth, suggesting that SlPP2C.Ds inhibit the function of SlLHA4. Importantly, this inhibition was alleviated by the addition of SlSAURs to the SlPP2C.Ds-SlLHA4 system, indicating that SlSAURs can counteract the inhibitory effects of SlPP2C.Ds on SlLHA4, thereby promoting PM H^+^-ATPase activity.

**Figure 5.**
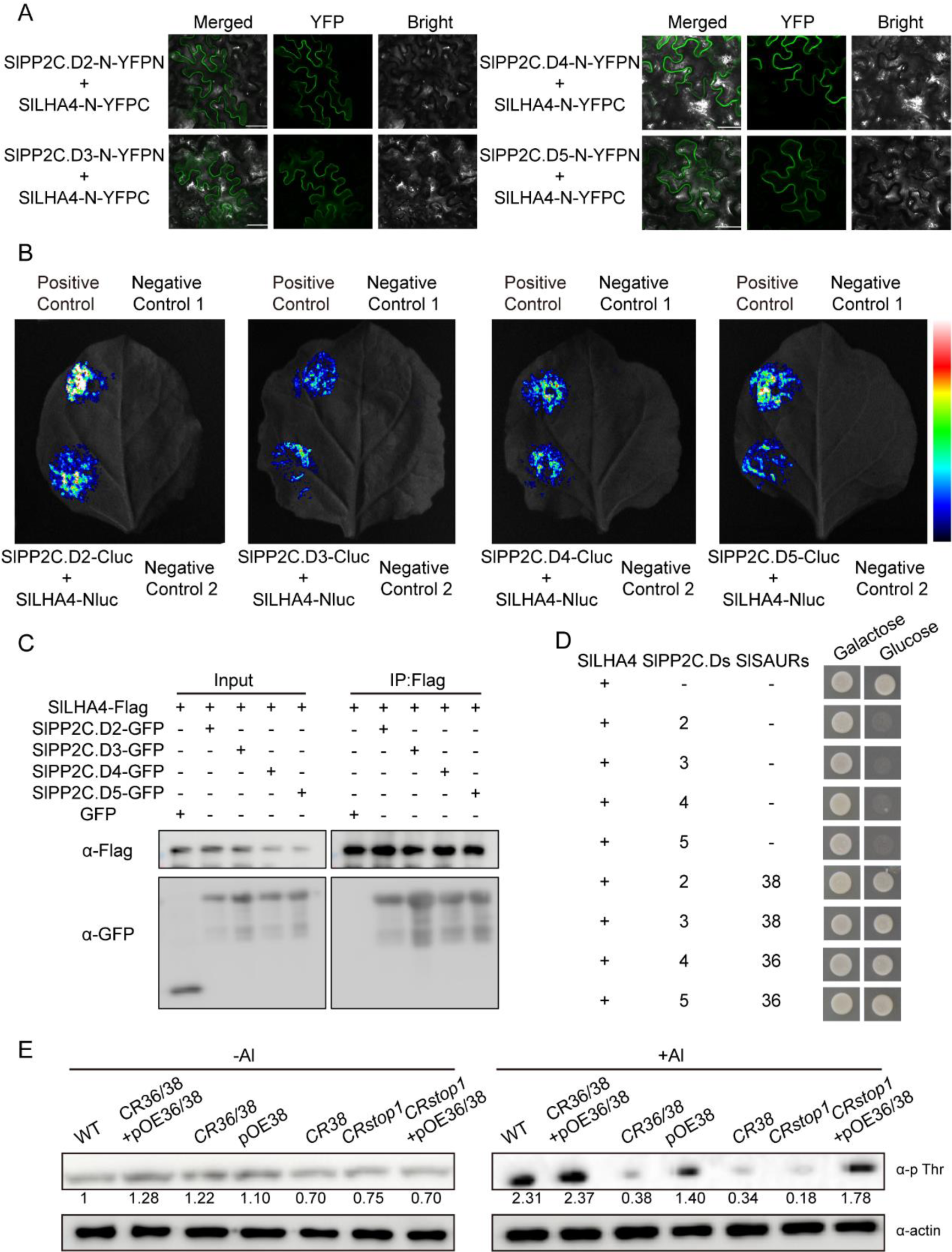
SlSAURs promote PM H^+^-ATPase activity by inhibiting SlPP2C.Ds phosphatase activity. **A** BiFC assay. SlPP2C.Ds-N-YFPN and SlLHA4-N-YFPC were transiently expressed in tobacco leaves. Scale bar =50 μm. **B** LCI assay. The paired constructs were co-infiltrated in *N. benthamiana* leaves, after 4-day incubation, luciferase was detected. Positive control was SlSTOP1-Nluc and SlSZP1-Cluc. Negative control 1 represented SlPP2C.Ds-Cluc co-infiltrated with SlSTOP1-Nluc. Negative control 2 represented SlLHA4-Nluc co-infiltrated with SlSZP1-Cluc. **C** Co-IP assay. SlPP2C.Ds-GFP was co-infiltrated with SlLHA4-Flag in N. *benthamiana* leaves. After 4 days, total protein extracts were immunoprecipitated with anti-Flag agarose beads and then detected with anti-GFP antibody. **D** SlLHA4 complementation of PM H^+^-ATPase activity in yeast RS72. Full length of *SlSAURs*, *SlPP2C.Ds* and *SlLHA4* were cloned into pMP1612, pMP1645 and pMP1745, respectively. The paired constructs were transformed into RS72 and then grown on glucose (pH 6.5) selective medium. **E** Phosphorylation level of PM H^+^-ATPase under normal condition and aluminum stress. Purified SlLHA4-GST were co-incubated with total protein of different materials about 30 minutes. The purified SlLHA4 with total protein was co-incubated with anti-GST agarose beads and the phosphorylation level of SlLHA4 was detected by anti-pThr antibody. Anti-actin antibody was used to detect the total protein level.

We also examined the post-translational modification of PM H^+^-ATPase by phosphorylation, which is crucial for its activation. Using a phosphorylation-specific antibody, we assessed the effect of aluminum stress on PM H*^+^*-ATPase phosphorylation levels (Figure 5E). The purified SlLHA4-GST was co-incubated with plant total protein, and under normal condition, all lines showed similar phosphorylation level. However, following aluminum treatment, an increase in phosphorylation was observed in WT, while knock out lines showed lower phosphorylation levels. Notably, complementation with *SlSAUR* restored phosphorylation levels compared to the *Slsaurs* or *Slstop1*.In summary, SlSAURs activate the activity of the plasma membrane proton pump by interacting with SlPP2C.D to release the inhibition of SlPP2C.D on SlLHA4.

## Discussion

In this study, we have elucidated a pivotal regulatory mechanism involving *SlSAUR36* and *SlSAUR38*, which functions as direct downstream targets of the key transcription factor SlSTOP1 and its enhancer SlSZP1, within the context of aluminum tolerance (Figure 1 and 2). Our research has shed light on the molecular underpinnings of the observed growth inhibition in root elongation under aluminum stress conditions, particularly when SlSTOP1 is non-functional. We have established that SlPP2C.D interacts with SlLHA4, a plasma membrane proton pump, to inhibit its activity, thereby affecting root elongation (Figure 5). While SlSAUR36/38 interact with SlPP2C.Ds (Figure 4), inhibiting their phosphatase activity and consequently releasing SlLHA4 to activate proton pumps and promote cell expansion (Figure 6). Collectively, these findings reveal a complex interplay between transcriptional regulation and post-translational modification that is central to the plant’s adaptive response to aluminum stress, highlighting the significance of SlSAUR36/38 in modulating root growth and development within this context.

**Figure 6.**
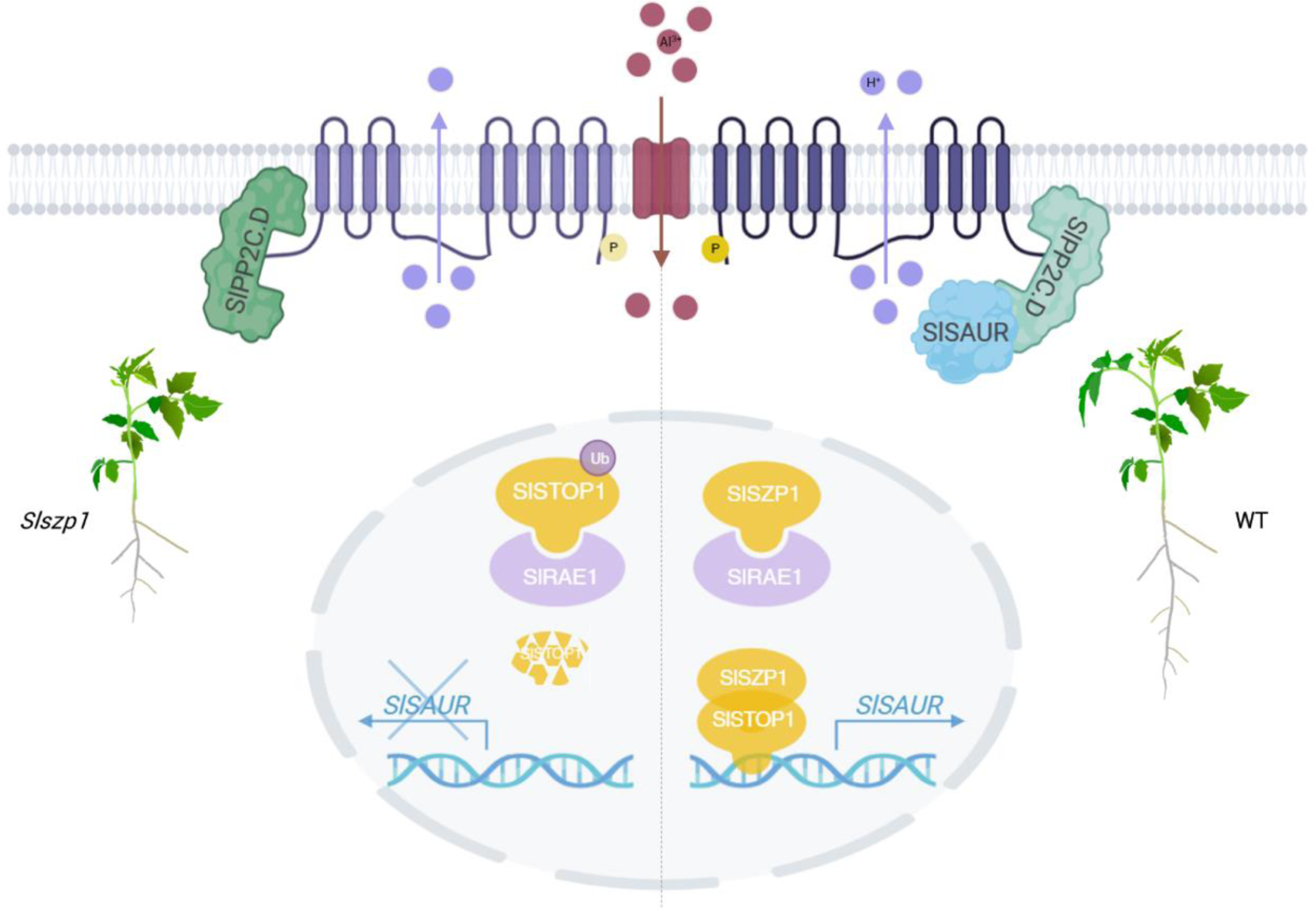
Model for the SlSAURs-SlPP2C.D-SlLHA under aluminum stress. When Al^3+^ is detected by plant cell, the transcription factor SlSTOP1 accumulates rapidly, and forms a complex with SlSZP1enhancer. They then bind to the promoter of SlSAURs to active their expression. The SlSAUR proteins, interact with the SlPP2C.D phosphatases, leading to the inhibition of the phosphatase activity. This inhibition releases the inhibition of SlPP2C.D on SlLHA4, a plasma membrane H^+^-ATPase. With reduced phosphatase activity, SlLHA4 remains phosphorylated, which in turn enhances the proton pump activity, promoting cell expansion and resulting in elongated root growth under conditions of aluminum stress.

Under aluminum stress, roots can secrete a variety of organic acids, such as malic acid, citric acid, and oxalate acid, which play a crucial role in chelating aluminum ions. This process, alongside the sequestration of aluminum ions within cellular compartments, serves as a strategic response to alleviate the damage of aluminum ions to roots. Notably, ALMT1 (Al-activated malate transporter 1) and MATE (Multidrug and Toxic Compound Extrusion) transporters which are downstream targets of STOP1, are known to facilitate aluminum tolerance by releasing malate and citrate, respectively, to chelate aluminum ions (Hoekenga et al., 2006; Iuchi et al., 2007; Liu et al., 2009). OsALS1, an ABC transporter-like protein, transfer aluminum ions from root tip to vacuole to avoid aluminum damage (Huang et al., 2012). At present, the research on target genes of STOP1 has predominantly centered on understanding how plants avoid or tolerate aluminum ions. However, the most significant damage caused by aluminum stress is the substantial inhibition of root length, a phenomenon that is not well understood and has received less attention.

Historically, it has been established that aluminum impacts the cell expansion within the root (Ryan et al., 1993; Gunse et al., 1997; Doncheva et al., 2005). The inhibition of root by aluminum toxics was primarily attributed to the reduction in cell expansion rather than a decrease in cell division (Doncheva et al., 2005). This raises the question of the precise manner in which aluminum impacts cell expansion. In our research, we identified *SlSAUR36* and *SlSAUR38* as novel targets of SlSTOP1. The SAUR family, known for its significant role in the early response to auxin and being the largest such family, has increasingly been recognized for its crucial regulatory functions in plant growth and development (Chae et al., 2012; Wang et al., 2020; Cui et al., 2023) since its initial discovery (Mcclure and Guilfoyle, 1987). This led us to hypothesize that SAURs could be the key factors in the reduction of root cell expansion under aluminum stress.

In previous research, we found that in normal condition, *Slstop1* and wild type showed no difference in root length, suggesting that SlSTOP1 does not regulate root length in the absence of aluminum stress (Zhang et al., 2022). However, when aluminum ions stimulated plant roots, it caused accumulation of SlSTOP1, thereby increasing the expression of downstream genes, exhibiting significant differences in root length (Zhang et al., 2022). In the current study, we found that under normal conditions, the root lengths of lines overexpressing *SlSAUR36* or *SlSAUR38*, as well as the *Slsaur36* and *Slsaur38* mutants, did not differ from that of *Slstop1* and WT (Figure 3). When treated with aluminum ions, the *Slsaur36* and *Slsaur38* exhibited shorter root length than wild type, whereas the overexpression lines showed longer root length. This further confirmed that reduced root elongation in *Slstop1* was attributed to the absence of SlSTOP1 protein, which consequently results in a failure to upregulate *SlSAURs*. Under aluminum stress, accumulated SlSTOP1 could arouse the expression of *SlSAUR36* and *SlSAUR38* to promote root elongation.

Our results also highlight the potential functional redundancy within the large *SAUR* gene family in tomato. With 99 SlSAUR members identified, it is plausible that the simultaneous knockout of *SlSAUR36* and *SlSAUR38* does not result in root developmental defects due to the compensatory roles of other family members. This redundancy may ensure normal organ development and growth under various stress conditions. STOP1, a transcription factor known to respond to aluminum ions, has also been implicated in other stress responses, including drought, salt, and low phosphate stress (Balzergue et al., 2017; Sadhukhan et al., 2019). Under aluminum stress, accumulated STOP1 promoted the efflux of malic acid through ALMT (Iuchi et al., 2007). At low Pi/Fe ratio, STOP1 significantly accumulated in the nucleus, promoting *ALMT* expression, enhancing root malic acid efflux (Balzergue et al., 2017). The STOP1-ALMT model could be used to many stresses, because of the multifunction of malate. This suggest SlSTOP1-SlSAUR model could potentially explain the root length inhibition pathway under other stress mediated by STOP1. OsSAUR39 has been shown to respond to transient changes in nitrogen conditions, salinity stress, and anoxia (Walia et al., 2005; Lasanthi-Kudahettige et al., 2007; Kant et al., 2009). STOP1 can also accumulated in salinity stress (Sadhukhan et al., 2019), and responded to low-oxygen stress (Enomoto et al., 2019). In these STOP1 regulated root phenotypes, SAUR, which can also be induced, in regulating root length under stress, presents an intriguing question for future research.

Auxin plays a key role in root development in response to Al stress. In maize, aluminum reduced the accumulation of auxin in root tip by inducing the expression of *ZmPGP1*, leading to shorter root length (Zhang et al., 2018). The addition of indole-3-acetic acid (IAA) to the elongation zone (EZ) significantly alleviated Al-induced root growth inhibition in maize (Kollmeier et al., 2000). In soybean, auxin increased the exudation of citrate by enhancing the expression of *MATE* and increased the phosphorylation of plasma membrane H^+^-ATPase (Wang et al., 2016). Al^3+^induced the expression of *CsUGT84J2* and regulated endogenous auxin homeostasis in tea roots, to promote the growth of tea (Jiang et al., 2023). The effect of IAA on aluminum stress varied among plant species. The specific effect of IAA on tomato root under aluminum stress remains unclear. After applying IAA, *SlSAUR36\38* were up-regulated in five minutes (Wu et al., 2012), suggesting their potential involvement in the auxin signaling pathway. SAUR15 has been shown to promote root accumulation and lateral root development (Yin et al., 2020). The connection of auxin signal pathway and aluminum stress in tomato needs further exploration.

PM H^+^-ATPases, which generate proton gradients and electrical potential differences (Palmgren, 2001; Falhof et al., 2016), are crucial for various physiological functions. Environmental factors can change the expression levels of PM H^+^-ATPase genes (Yang et al., 2011; Yuan et al., 2017). Post-translational modification of PM H^+^-ATPase by phosphorylation and dephosphorylation significantly impacts its activity. Abiotic stress, hormones, fungi toxins can induce the phosphorylation or dephosphorylation of the PM H^+^-ATPase (Yang et al., 2019; Miao et al., 2021). More and more evidence suggest that proton pumps are also involved in aluminum stress response. In squash (*Cucurbita pepo L. cv Tetsukabuto*), aluminum reduced the activity of PM H^+^-ATPase significantly (Ahn et al., 2001). In contrast, in tomato, aluminum induced the expression level of PM H^+^-ATPase genes (Yang et al., 2011). The activity of PM H^+^-ATPase was usually related to certain transporters. SAL1 (SENSITIVE TO ALUMINUM 1), a PP2C.D phosphatase, could interact with OSA (PM H^+^-ATPase) in rice, inhibiting its activity, and reducing the Al uptake by NRAT1 (Nramp aluminum transporter 1). In Arabidopsis, the activity of PM H^+^-ATPase was inhibited by PP2C.D5/6/7, and increased activity promoted the secretion of malate (Xie et al., 2023). Our findings that SlPP2C.D interacted with SlLHA4 to inhibit its activity (Figure 5) and that SlSAUR36/38 can counteract this inhibition (Figure 6), providing a model for maintaining H^+^-ATPase activity under aluminum stress. However, the initial phosphorylation event that activates H^+^-ATPase under aluminum stress remains to be elucidated. Transmembrane Kinase (TMK1), a key regulator in auxin signaling, directly interact with PM H^+^-ATPase within seconds, inducing phosphorylation to promote cell-wall acidification (Lin et al., 2021), may be a candidate for this activation.

SAUR proteins are typically involved in a range of physiological activities and are regulated by numerous transcription factors (TFs) (Franklin et al., 2011). SZP1, a potential C2H2-type transcription factor, can response to aluminum transcriptionally under aluminum stress (Zhang et al., 2022). In our research, we found not only SlSTOP1 can regulate *SlSAUR36* and *SlSAUR38*, but SlSZP1 also exerts regulatory control over these genes (Figure 2). Co-expressing SlSTOP1 and SlSZP1 may work together on the promoters of *SlSAUR36* and *SlSAUR38*, increasing their expression and promoting cell expansion under aluminum stress (Figure 2). On one hand, SlSTOP1, which undergoes post-translational modifications under aluminum stress, rapidly accumulates and promotes the expression of *SlSAURs*. On the other hand, SlSZP1, responding to aluminum signals significantly at the transcriptional level, leading to the rapid accumulation of SlSZP1 protein. The accumulated SlSZP1 not only prevents the degradation of SlSTOP1, but also promotes the gene expression of *SlSAURs*.

## Materials and methods

### Plant materials and growth conditions

In this study, we utilized two cultivars of *Solanum lycopersicum*: cv. Ailsa Craig (AC) and cv. Micro-Tom. All plant materials were cultivated in a greenhouse under natural light condition at China Agricultural University, Beijing. The aluminum treatment protocol was conducted in accordance with the methodology described in previous research (Zhang et al., 2022). All of the knock out lines confirmed to be homozygous and Cas9-free.

For *Slsaur36* single knock out line, *Slsaur36-1* line had a 1 bp deletion in target 2. For *Slsaur38* single knock out line, *Slsaur38-1* line had a 1 bp deletion in target 1. For *Slsaur36* and *Slsaur38* double knock out line, *Slsaur-168* has a 16 bp deletion in target 2 of *SlSAUR36* and 8 bp deletion in target 1 of *SlSAUR38*. *Slsaur-11* has a 1 bp deletion of *SlSAUR36* in target 2 and 1 bp deletion of *SlSAUR38* in target 1.

For *SlSAUR36* and *SlSAUR38* overexpressing lines and complementation lines, the coding sequence of *SlSAUR36* (351 bp) and *SlSAUR38* (474bp), excluding the stop codon, were driven by their native promoters were cloned into pCAMBIA-1305 using *EcoR* I and *Hind* III restriction sites. This process resulted in the following constructs: SlSAUR36p::SlSAUR36-1 (pOE36-1), SlSAUR36p::SlSAUR36-7/*Slstop1* (pOE36-7/*Slstop1*), SlSAUR38p::SlSAUR38-3 (pOE38-3), SlSAUR38p::SlSAUR38-6/*Slstop1* (pOE38-6/*Slstop1*) and SlSAUR36p::SlSAUR36-7+ SlSAUR38p::SlSAUR38-6/*Slstop1* (pOE36-7+ pOE38-6/*Slstop1*).

### RNA extraction and quantitative real-time PCR

The root samples treated and untreated with Al^3+^ were ground using liquid nitrogen. Total RNA was extracted with Trizol reagent. The Prime Script ^TM^ RT reagent Kit (TAKARA) was used to synthesize the first-strand cDNA. Quantitative Real-time PCR (qRT-PCR) was conducted on a 7500 Real-time PCR system. The expression analysis was performed with the 2^−△△Ct^ method. The primers were listed in Table S1. Three biological replicates were performed for each sample.

### Yeast one-hybrid (Y1H) assay

The full length CDS of *SlSTOP1* and *SlSZP1* were cloned into pGAD424 vector to produce SlSTOP1-pGAD424 and SlSZP1-pGAD424. The promoter fragments and the mutated fragments of *SlSAUR36* and *SlSAUR38*, as shown in Figure 1 and 2, were respectively constructed into the pLaczi vector. The SlSTOP1-pGAD424, SlSZP1-pGAD424 and the pLaczi vectors containing the respective fragments were co-transferred into YM4271 yeast strain (Weidi, Shanghai). As a negative control, the pGAD424 and fragment-containing pLaczi vectors were also co-transferred into YM4271 yeast strain. To detect protein-DNA interactions, a β-galactosidase assay was conducted using Z-buffer supplemented with X-gal (Fuxman Bass et al., 2016).

### Dual-luciferase assay

The full length CDS of *SlSTOP1* and *SlSZP1* were cloned into pGreen-62SK vector to produce SlSTOP1-62SK and SlSZP1-62SK. The promoter fragments of downstream genes, *SlSAUR36* and *SlSAUR38*, were constructed into the pGreen0800-mini-luciferase vector. These vectors were co-injected into *Agrobacterium* strain GV3101-P19-psoup (Weidi, Shanghai) according to the following combinations: SlSTOP1-62SK with *SlSAUR36* promoter, SlSTOP1-62SK with *SlSAUR38* promoter, SlSZP1-62SK with *SlSAUR36* promoter, SlSZP1-62SK with *SlSAUR38* promoter, SlSZP1-62SK with SlSTOP1-62SK and *SlSAUR36* promoter, and SlSZP1-62SK with SlSTOP1-62SK and *SlSAUR38* promoter. The negative controls consisted of the pGreen-62SK empty vector and *SlSAUR* promoter constructs, respectively. Following a 3-day incubation, they were infiltrated into tobacco (*Nicotiana benthamiana*) leaves. After 4 days, the luminescence of tobacco leaves was detected by a luminescence detection kit (Vazyme).

### Electrophoretic mobility shift assay (EMSA)

The full-length sequence of SlSTOP1 was cloned into the pGEX-4T-1vector to produce a SlSTOP1-4T-1 vector. The vector was transformed into *E. coli* BL21 for protein expression and purification. The 50-bp promoter fragments which were shown in Figure 1 from *SlSAUR36* (p-1) and *SlSAUR38* (p-1) were labeled by biotin. The sequence was shown in Table S1. The EMSA was performed to assess the protein-DNA interactions with an EMSA kit (Thermo Fisher Scientific), following the manufacturer’s guidelines to ensure standardization and reproducibility of the assay.

### Subcellular localization assay

The coding sequence of *SlSAURs*, excluding the stop codon, were cloned into pSuper1300-GFP vector, and one more GFP were added at the C terminus of GFP. PMH (Plasma Membrane-Localized H^+^-ATPase) was used as plasma membrane marker. The coding sequence of *SlPP2C.Ds*, excluding the stop codon, were cloned into pSuper1300-GFP vector. After infiltrating into *N. benthamiana* leaves, GFP fluorescence was observed after 4-days incubation by a confocal laser scanning microscopy (Leica SP8).

### Yeast two-hybrid (Y2H) assay

Y2H was conducted as previous research described (Wang et al., 2022).The full-length sequence of *SlSAUR36* and *SlSAUR38* were cloned into the pGBKT7 vector. And full-length sequence of *SlPP2C.Ds* were cloned into pGADT7 vector respectively. Two paired constructs were co-transformed into Y2H gold (Weidi, Shanghai). Finally, the yeast clones were selected and cultured in SD-TLHA supplemented with X-α-gal medium. Positive control group was SlSTOP1-BD and SlSZP1-AD, as previously described (Zhang et al., 2022).

### Bimolecular fluorescence complementation (BiFC)

Full-length of *SlSAUR36*, *SlSAUR38* were cloned into the pCAMBIA1300-35S-N-YFPN, and full-length of *SlPP2C.D2/3/4/5* were cloned into the pCAMBIA1300-35S-N-YFPC. Full-length of *SlPP2C.D2/3/4/5* were cloned into the pCAMBIA1300-35S-N-YFPN, and full-length of *SlLHA4* were cloned into the pCAMBIA1300-35S-N-YFPC.The combinations were respectively infiltrated into *N. benthamiana* leaves. After 4-day incubation, YFP fluorescence was observed by a confocal laser scanning microscopy (Leica SP8).

### Co-immunoprecipitation (Co-IP) assay

For the Co-IP assays, *N. benthamiana* leaves infiltrated with different combinations of constructs transformed *Agrobacterium* were ground with liquid nitrogen, and total protein was extracted by extraction buffer (20 mM Tris-HCl, pH 7.4, 1mM EDTA, 100mM NaCl, 1% NP-40, 1× protease inhibitor cocktail). After 10000r centrifugation under 4 ℃, the resulting supernatant was collected as the source of total protein. The supernatant was then incubated with anti-Flag agarose beads (SMART, Anti-DYKDDDDK Affinity Beads, SA042005) for 2 h at 4 ℃. Following the incubation, the beads were washed 3 times with extraction buffer. Finally, 40 μl elution buffer was added to beads to recover the immunoprecipitated proteins. The eluted protein samples were then resolved by sodium dodecyl sulfate-polyacrylamide gel electrophoresis (SDS-PAGE) and subjected to Western blot detection using anti-Flag and anti-GFP antibodies (Abcam, Cambridge, UK).

### Phosphatase activity assay

Phosphatase assay was performed according to previous reported research (Spartz et al., 2014). The full-length cDNA of *SlSAURs* and *SlPP2C.Ds* were cloned into pMAL-p5x and pGEX-4T-1, respectively. The reaction system contains 75 mM Tris (pH 7.6), 10 mM MnCl_2_, 100 mM NaCl, 0.5 mM EDTA, and 5 mM pNPP, with the final reaction volume was adjusted to 100 μl. The reaction was initiated by the addition of the enzyme, and the absorbance at 405 nm was continuously monitored for a period of 10 minutes using a microplate reader (MD i3x) to measure the release of p-nitrophenol from pNPP, indicative of phosphatase activity.

### Yeast complementation assay

*S. cerevisiae* strain *RS72* was used for complementation tests (Fuglsang et al., 2007). The full length of *SlLHA4*, *SlPP2C.D* and *SlSAURs* were respectively cloned into pMP1745, pMP1645 and pMP1612. The complementation assay was carried out following the methodology described in (Yang et al., 2019), which allows for the functional analysis of the cloned genes within the yeast strain.

### Molecular docking

To elucidate the three-dimensional structure of the protein of interest, the corresponding amino acid sequence was submitted to the SWISS-MODEL server (https://swissmodel.expasy.org/), a repository for protein 3D structure homology modeling. Upon completion of the homology modeling, the predicted protein structure was downloaded in Protein Data Bank (PDB) format. The PDB file was subsequently uploaded to the ZDOCK server (https://zdock.umassmed.edu/) for protein docking analysis. The resulting PDB files from the docking analysis were then imported into the PDBe website (https://www.ebi.ac.uk/pdbe/) for comprehensive analysis. Finally, the docking result with the highest score, indicating the most favorable interaction, was selected and imported into PyMOL molecular visualization software (https://pymol.org/).

### Statistical analyses

Data were analyzed by ANOVA with Tukey’s multiple comparison test or t-test by GraphPad Prism 10.

### Accession numbers

Sequence data from this article can be found in the Solanaceae Genomics Network (https://solgenomics.net/). SlSAUR36 (Solyc03g082520), SlSAUR38 (Solyc03g097510), SlPP2C.D2 (Solyc02g083420), SlPP2C.D3 (Solyc03g033340), SlPP2C.D4 (Solyc05g055980), SlPP2C.D5 (Solyc06g065920), SlLHA4 (Solyc07g017780).

## Acknowledgement

We thank Professor Yan Guo (College of Biological Sciences, China Agricultural University) and Professor Yongqing Yang (College of Biological Sciences, China Agricultural University) for providing yeast strain *RS72* and pMP1745, pMP1645 and pMP1612 vectors.

## Author Contributions

Y-DG, NZ and DD designed this project. DD, QD, JZ, CJ performed the experiments. DD, QD, LZ and HL analyzed the data. DD, NZ and Y-DG wrote the manuscript. All authors read and approved the final manuscript. DD and QD contributed equally to this work.

## Reference

Agrahari RK, Enomoto T, Ito H, Nakano Y, Yanase E, Watanabe T, Sadhukhan A, Iuchi S, Kobayashi M, Panda SK, Yamamoto YY, Koyama H, Kobayashi Y (2021) Expression GWAS of *PGIP1* Identifies STOP1-Dependent and STOP1-Independent Regulation of PGIP1 in Aluminum Stress Signaling in *Arabidopsis*. Front. Plant Sci. 12

Ahn SJ, Sivaguru M, Osawa H, Chung GC, Matsumoto H (2001) Aluminum Inhibits the H^+^-ATPase Activity by Permanently Altering the Plasma Membrane Surface Potentials in Squash Roots1. Plant Physiol. 126: 1381–1390

Balzergue C, Dartevelle T, Godon C, Laugier E, Meisrimler C, Teulon JM, Creff A, Bissler M, Brouchoud C, Hagège A, Muller J, Chiarenza S, Javot H, Becuwe-Linka N, David P, Péret B, Delannoy E, Thibaud MC, Armengaud J, Abel S, Pellequer JL, Nussaume L, Desnos T (2017) Low phosphate activates STOP1-ALMT1 to rapidly inhibit root cell elongation. Nat. Commun. 8: 15300

Chae K, Isaacs CG, Reeves PH, Maloney GS, Muday GK, Nagpal P, Reed JW (2012) *Arabidopsis SMALL AUXINUPRNA63* promotes hypocotyl and stamen filament elongation. Plant J. 71: 684–697

Cui M, Li Y, Li J, Yin F, Chen X, Qin L, Wei L, Xia G, Liu S (2023) Ca^2+^-dependent TaCCD1 cooperates with TaSAUR215 to enhance plasma membrane H^+^-ATPase activity and alkali stress tolerance by inhibiting PP2C-mediated dephosphorylation of TaHA2 in wheat. Mol. Plant. 16: 571–587

Doncheva S, Amenós M, Poschenrieder C, Barceló J (2005) Root cell patterning: a primary target for aluminum toxicity in maize. J. Exp. Bot. 56: 1213–1220

Dong J, Sun N, Yang J, Deng ZG, Lan JQ, Qin GJ, He H, Deng XW, Irish VF, Chen HD, Wei N (2019) The Transcription Factors TCP4 and PIF3 Antagonistically Regulate Organ-Specific Light Induction of *SAUR* Genes to Modulate Cotyledon Opening during De-Etiolation in Arabidopsis. Plant Cell 31: 1155–1170

Enomoto T, Tokizawa M, Ito H, Iuchi S, Kobayashi M, Yamamoto YY, Kobayashi Y, Koyama H (2019) STOP1 regulates the expression of *HsfA2* and *GDHs* that are critical for low-oxygen tolerance in Arabidopsis. J. Exp. Bot. 70: 3297–3311

Falhof J, Pedersen JT, Fuglsang AT, Palmgren M (2016) Plasma Membrane H^+^-ATPase Regulation in the Center of Plant Physiology. Mol. Plant. 9: 323–337

Fendrych M, Leung J, Friml J (2016) TIR1/AFB-Aux/IAA auxin perception mediates rapid cell wall acidification and growth of Arabidopsis hypocotyls. eLife 5

Franklin KA, Lee SH, Patel D, Kumar SV, Spartz AK, Gu C, Ye SQ, Yu P, Breen G, Cohen JD, Wigge PA, Gray WM (2011) PHYTOCHROME-INTERACTING FACTOR 4 (PIF4) regulates auxin biosynthesis at high temperature. Proc. Natl. Acad. Sci. U. S. A. 108: 20231–20235

Fuglsang AT, Guo Y, Cuin TA, Qiu QS, Song CP, Kristiansen KA, Bych K, Schulz A, Shabala S, Schumaker KS, Palmgren MG, Zhu JK (2007) *Arabidopsis* Protein Kinase PKS5 Inhibits the Plasma Membrane H^+^-ATPase by Preventing Interaction with 14-3-3 Protein. Plant Cell 19: 1617–1634

Fuxman Bass JI, Reece-Hoyes JS, Walhout AJM (2016) Colony Lift Colorimetric Assay for β-Galactosidase Activity. Cold Spring Harb. Protoc. 2016: pdb.prot088963

Gunse B, Poschenrieder C, Barcelo J (1997) Water transport properties of roots and root cortical cells in proton-and Al-stressed maize varieties. Plant Physiol. 113: 595–602

Guo JH, Liu XJ, Zhang Y, Shen JL, Han WX, Zhang WF, Christie P, Goulding KWT, Vitousek PM, Zhang FS (2010) Significant Acidification in Major Chinese Croplands. Science 327: 1008–1010

He YJ, Liu Y, Li MZ, Lamin-Samu AT, Yang DD, Yu XL, Izhar M, Jan I, Ali M, Lu G (2021) The Arabidopsis SMALL AUXIN UP RNA32 Protein Regulates ABA-Mediated Responses to Drought Stress. Front. Plant Sci. 12: 625493

Hoekenga OA, Maron LG, Piñeros MA, Cançado GMA, Shaff J, Kobayashi Y, Ryan PR, Dong B, Delhaize E, Sasaki T, Matsumoto H, Yamamoto Y, Koyama H, Kochian LV (2006) *AtALMT1*, which encodes a malate transporter, is identified as one of several genes critical for aluminum tolerance in *Arabidopsis*. Proc. Natl. Acad. Sci. U. S. A. 103: 9738–9743

Huang CF, Yamaji N, Chen ZC, Ma JF (2012) A tonoplast-localized half-size ABC transporter is required for internal detoxification of aluminum in rice. Plant J. 69: 857–867

Iuchi S, Koyama H, Iuchi A, Kobayashi Y, Kitabayashi S, Kobayashi Y, Ikka T, Hirayama T, Shinozaki K, Kobayashi M (2007) Zinc finger protein STOP1 is critical for proton tolerance in *Arabidopsis* and coregulates a key gene in aluminum tolerance. Proc. Natl. Acad. Sci. U. S. A. 104: 9900–9905

Jiang XL, Lai SY, Kong DX, Hou XH, Shi YF, Fu ZP, Liu YJ, Gao LP, Xia T (2023) Al-induced CsUGT84J2 enhances flavonol and auxin accumulation to promote root growth in tea plants. Hort. Res 10

Kant S, Bi YM, Zhu T, Rothstein SJ (2009) *SAUR39*, a Small Auxin-Up RNA Gene, Acts as a Negative Regulator of Auxin Synthesis and Transport in Rice. Plant Physiol. 151: 691–701

Kodaira KS, Qin F, Tran LSP, Maruyama K, Kidokoro S, Fujita Y, Shinozaki K, Yamaguchi-Shinozaki K (2011) Arabidopsis Cys2/His2 Zinc-Finger Proteins AZF1 and AZF2 Negatively Regulate Abscisic Acid-Repressive and Auxin-Inducible Genes under Abiotic Stress Conditions. Plant Physiol. 157: 742–756

Kollmeier M, Felle HH, Horst WJ (2000) Genotypical Differences in Aluminum Resistance of Maize Are Expressed in the Distal Part of the Transition Zone. Is Reduced Basipetal Auxin Flow Involved in Inhibition of Root Elongation by Aluminum? Plant Physiol. 122: 945–956

Lasanthi-Kudahettige R, Magneschi L, Loreti E, Gonzali S, Licausi F, Novi G, Beretta O, Vitulli F, Alpi A, Perata P (2007) Transcript Profiling of the Anoxic Rice Coleoptile. Plant Physiol. 144: 218–231

Lin WW, Zhou X, Tang WX, Takahashi K, Pan X, Dai JW, Ren H, Zhu XY, Pan SQ, Zheng HY, Gray WM, Xu TD, Kinoshita T, Yang ZB (2021) TMK-based cell-surface auxin signalling activates cell-wall acidification. Nature 599: 278–282

Liu JP, Magalhaes JV, Shaff J, Kochian LV (2009) Aluminum-activated citrate and malate transporters from the MATE and ALMT families function independently to confer Arabidopsis aluminum tolerance. Plant J. 57: 389–399

Llugany M, Poschenrieder C, Barceló J (2008) Monitoring of aluminum-induced inhibition of root elongation in four maize cultivars differing in tolerance to aluminum and proton toxicity. Physiol. Plant. 93: 265–271

Ma PA, Chen X, Liu C, Meng YH, Xia ZQ, Zeng CY, Lu C, Wang WQ (2017) MeSAUR1, Encoded by a Small Auxin-Up RNA Gene, Acts as a Transcription Regulator to Positively Regulate ADP-Glucose Pyrophosphorylase Small Subunit1a Gene in Cassava. Front. Plant Sci. 8: 1315

Marathe S, Grotewold E, Otegui MS (2024) Should I stay or should I go? Trafficking of plant extra-nuclear transcription factors. Plant Cell 36: 1524–1539

Mcclure BA, Guilfoyle T (1987) Characterization of a class of small auxin-inducible soybean polyadenylated RNAs. Plant Mol.Biol. 9: 611–623

Miao R, Yuan W, Wang Y, García Maquilón I, Dang X, Li Y, Zhang J, Zhu Y, Rodriguez P, Xu W (2021) Low ABA concentration promotes root growth and hydrotropism through relief of ABA INSENSITIVE 1-mediated inhibition of plasma membrane H^+^-ATPase 2. Sci. Adv. 7: eabd4113

Młodzińska E, Kłobus G, Christensen MD, Fuglsang AT (2015) The plasma membrane H^+^-ATPase AHA2 contributes to the root architecture in response to different nitrogen supply. Physiol. Plant. 154: 270–282

Oh E, Zhu JY, Bai MY, Arenhart RA, Sun Y, Wang ZY (2014) Cell elongation is regulated through a central circuit of interacting transcription factors in the Arabidopsis hypocotyl. eLife 3: e03031

Ownby JD, Popham HR (1990) Citrate Reverses the Inhibition of Wheat Root Growth Caused by Aluminum. J. Plant Physiol. 135: 588–591

Palmgren MG (2001) PLANT PLASMA MEMBRANE H^+^-ATPases: Powerhouses for Nutrient Uptake. Annu. Rev. Plant Physiol. 52: 817–845

Ren H, Park MY, Spartz AK, Wong JH, Gray WM (2018) A subset of plasma membrane-localized PP2C.D phosphatases negatively regulate SAUR-mediated cell expansion in Arabidopsis. PLoS Genet. 14: e1007455

Ryan PR, Ditomaso JM, Kochian LV (1993) Aluminum Toxicity in Roots: An Investigation of Spatial Sensitivity and the Role of the Root Cap. J. Exp. Bot. 44: 437–446

Sadhukhan A, Enomoto T, Kobayashi Y, Watanabe T, Iuchi S, Kobayashi M, Sahoo L, Yamamoto YY, Koyama H (2019) Sensitive to Proton Rhizotoxicity1 Regulates Salt and Drought Tolerance of *Arabidopsis thaliana* through Transcriptional Regulation of *CIPK23*. Plant Cell Physiol. 60: 2113–2126

Sadhukhan A, Kobayashi Y, Iuchi S, Koyama H (2021) Synergistic and antagonistic pleiotropy of STOP1 in stress tolerance. Trends Plant Sci. 26: 1014–1022

Sawaki Y, Iuchi S, Kobayashi Y, Kobayashi Y, Ikka T, Sakurai N, Fujita M, Shinozaki K, Shibata D, Kobayashi M, Koyama H (2009) STOP1 Regulates Multiple Genes That Protect Arabidopsis from Proton and Aluminum Toxicities. Plant Physiol. 150: 281–294

Spartz AK, Lee SH, Wenger JP, Gonzalez N, Itoh H, Inzé D, Peer WA, Murphy AS, Overvoorde PJ, Gray WM (2012) The SAUR19 subfamily of *SMALL AUXIN UP RNA* genes promote cell expansion. Plant J. 70: 978–990

Spartz AK, Ren H, Park MY, Grandt KN, Lee SH, Murphy AS, Sussman MR, Overvoorde PJ, Gray WM (2014) SAUR Inhibition of PP2C-D Phosphatases Activates Plasma Membrane H+-ATPases to Promote Cell Expansion in *Arabidopsis*. Plant Cell 26: 2129–2142

Takahashi K, Hayashi K, Kinoshita T (2012) Auxin Activates the Plasma Membrane H^+^-ATPase by Phosphorylation during Hypocotyl Elongation in Arabidopsis. Plant Physiol. 159: 632–641

Tokizawa M, Enomoto T, Ito H, Wu LJ, Kobayashi Y, Mora-Macias J, Armenta-Medina D, Iuchi S, Kobayashi M, Nomoto M, Tada Y, Fujita M, Shinozaki K, Yamamoto YY, Kochian LV, Koyama H (2021) High affinity promoter binding of STOP1 is essential for early expression of novel aluminum-induced resistance genes *GDH1* and *GDH2* in Arabidopsis. J. Exp. Bot. 72: 2769–2789

Tokizawa M, Kobayashi Y, Saito T, Kobayashi M, Iuchi S, Nomoto M, Tada Y, Yamamoto YY, Koyama H (2015) SENSITIVE TO PROTON RHIZOTOXICITY1, CALMODULIN BINDING TRANSCRIPTION ACTIVATOR2, and other transcription factors are involved in ALUMINUM-ACTIVATED MALATE TRANSPORTER1 expression. Plant Physiol 167: 991-1003

Tsutsui T, Yamaji N, Ma JF (2011) Identification of a Cis-Acting Element of ART1, a C2H2-Type Zinc-Finger Transcription Factor for Aluminum Tolerance in Rice. Plant Physiol. 156: 925–931

Walia H, Wilson C, Condamine P, Liu X, Ismail AM, Zeng LH, Wanamaker SI, Mandal J, Xu J, Cui XP, Close TJ (2005) Comparative Transcriptional Profiling of Two Contrasting Rice Genotypes under Salinity Stress during the Vegetative Growth Stage. Plant Physiol. 139: 822–835

Wang JJ, Sun N, Zhang FF, Yu RB, Chen HD, Deng XW, Wei N (2020) SAUR17 and SAUR50 Differentially Regulate PP2C-D1 during Apical Hook Development and Cotyledon Opening in Arabidopsis. Plant Cell 32: 3792–3811

Wang M, Yang S, Sun L, Feng Z, Gao Y, Zhai X, Dong Y, Wu H, Cui Y, Li S, Xu S, Bartholomew ES, Ren H, Liu X (2022) A CBL4-CIPK6 module confers salt tolerance in cucumber. Vegetable Research 2: 1–10

Wang P, Yu WQ, Zhang JR, Rengel Z, Xu J, Han QQ, Chen LM, Li KZ, Yu YX, Chen Q (2016) Auxin enhances aluminum-induced citrate exudation through upregulation of *GmMATE* and activation of the plasma membrane H^+^-ATPase in soybean roots. Ann. Bot. 118: 933–940

Wang S, Wang Q, Jiang W, Wang Y, Yan J, Li X, Wang J, Guan Q, Ma F, Zhang J, Zheng Q, Zou Y, Xu J (2024) Evaluating the sustainable cultivation of ‘Fuji’ apples: suitable crop load and the impact of chemical thinning agents on fruit quality and transcription. Fruit Research 4

Wang XY, Yu RB, Wang JJ, Lin ZC, Han X, Deng ZG, Fan LM, He H, Deng XW, Chen HD (2020) The Asymmetric Expression of *SAUR* Genes Mediated by ARF7/19 Promotes the Gravitropism and Phototropism of Plant Hypocotyls. Cell Reports 31: 107529

Wong JH, Spartz AK, Park MY, Du MM, Gray WM (2019) Mutation of a Conserved Motif of PP2C.D Phosphatases Confers SAUR Immunity and Constitutive Activity. Plant Physiol. 181: 353–366

Wu J, Liu SY, He YJ, Guan XY, Zhu XF, Cheng L, Wang J, Lu G (2012) Genome-wide analysis of SAUR gene family in Solanaceae species. Gene 509: 38–50

Xie WX, Liu S, Gao HL, Wu J, Liu DL, Kinoshita T, Huang CF (2023) PP2C.D phosphatase SAL1 positively regulates aluminum resistance via restriction of aluminum uptake in rice. Plant Physiol. 192: 1498-1516

Yang JL, Zhu XF, Peng YX, Zheng C, Ming F, Zheng SJ (2011) Aluminum regulates oxalate secretion and plasma membrane H^+^-ATPase activity independently in tomato roots. Planta 234: 281–291

Yang Y, Wu Y, Ma L, Yang Z, Dong Q, Li Q, Ni X, Kudla J, Song C, Guo Y (2019) The Ca^2+^ Sensor SCaBP3/CBL7 Modulates Plasma Membrane H^+^-ATPase Activity and Promotes Alkali Tolerance in Arabidopsis. Plant Cell 31: 1367–1384

Yin HJ, Li MZ, Lv MH, Hepworth SR, Li DD, Ma CF, Li J, Wang SM (2020) SAUR15 Promotes Lateral and Adventitious Root Development via Activating H^+^-ATPases and Auxin Biosynthesis. Plant Physiol. 184: 837–851

Yuan W, Zhang DP, Song T, Xu FY, Lin S, Xu WF, Li QF, Zhu YY, Liang JS, Zhang JH (2017) Arabidopsis plasma membrane H^+^-ATPase genes *AHA2* and *AHA7* have distinct and overlapping roles in the modulation of root tip H^+^ efflux in response to low-phosphorus stress. J. Exp. Bot. 68: 1731–1741

Zhang F, Yan X, Han X, Tang R, Chu M, Yang Y, Yang Y-H, Zhao F, Fu A, Luan S, Lan W (2019) A Defective Vacuolar Proton Pump Enhances Aluminum Tolerance by Reducing Vacuole Sequestration of Organic Acids. Plant Physiol. 181: 743–761

Zhang L, Dong DH, Wang JF, Wang ZR, Zhang JJ, Bai RY, Wang XW, Wilhe i MDR, Blumwald E, Zhang N, Guo YD (2022) A zinc finger protein SlSZP1 protects SlSTOP1 from SlRAE1mediated degradation to modulate aluminum resistance. New Phytol. 236: 165–181

Zhang ML, Lu XD, Li CL, Zhang B, Zhang CY, Zhang XS, Ding ZJ (2018) Auxin Efflux Carrier ZmPGP1 Mediates Root Growth Inhibition under Aluminum Stress. Plant Physiol. 177: 819–832

